# The effects of dietary iron supplementation on bacterial infections in *Manduca sexta* larval hemolymph

**DOI:** 10.64898/2026.03.21.713330

**Authors:** Marilyn K. Reese, Michael R. Kanost, Maureen J. Gorman

## Abstract

Iron is an essential nutrient for all types of organisms, including insects and the microbes that infect them. We predicted that insects fed an iron-supplemented diet would accumulate more iron in their hemolymph, and, because infectious microbes acquire iron from their hosts, that this extra iron would increase the severity of bacterial infections. To test this hypothesis, we studied the effects of dietary iron supplementation on infection outcomes in *Manduca sexta* (tobacco hornworm). Larvae were fed an artificial diet, with or without antibiotics, or the same diets supplemented with 10 mM iron. Control and iron-treated larvae were inoculated with non-pathogenic *Escherichia coli* or the entomopathogenic *Enterococcus faecalis*, and bacterial load and larval survival were measured. We found that dietary iron supplementation increased the iron content of hemolymph by approximately 20 fold; however, contrary to our prediction, this increase in iron did not result in an increase in the bacterial load of either *E. coli* or *E. faecalis*. The effect of iron supplementation on survival was more complicated. As expected, for larvae inoculated with nonpathogenic *E. coli*, iron supplementation had no effect. For larvae inoculated with *E. faecalis*, the effect of iron supplementation depended on whether antibiotics were present in the diet. Without antibiotics, iron supplementation prolonged larval survival; with antibiotics, iron supplementation decreased larval survival. The results of this study do not support the hypothesis that dietary iron supplementation increases infection severity in *M. sexta*. Instead, the results support the viewpoint that the relationship between dietary iron and infection outcome is complex.

## Introduction

Iron is an essential nutrient for all types of organisms, including insects and the microbes that infect them (1,2). This need for iron establishes a competition between microbes and their hosts for iron. Infectious microbes have evolved mechanisms for obtaining iron from their hosts, while hosts have evolved mechanisms that limit the ability of the infectious microbes to take up iron from their hosts, thereby limiting the severity of infections (3–5).

We were interested in the question of what happens to infection severity when a host insect consumes extra iron. Previous research has shown that dietary iron supplementation increases the amount of iron in insect bodies (6–8); therefore, one possible outcome would be that the additional host iron promotes growth of infectious agents, resulting in more severe infections. The number of studies testing this hypothesis are still somewhat limited. *Drosophila melanogaster fed* an iron-supplemented diet were more likely than those fed a control diet to die from the fungal pathogen *Rhizopus oryzae*, whereas flies fed the same diet containing an iron chelator that decreases iron bioavailability were less likely to die from the fungal infection (9). In contrast, the addition of iron or iron chelator to the diet had no effect on the number of the intracellular bacterium *Spiroplasma poulsonii* in infected flies, and dietary iron supplementation in flies did not increase the density of the intracellular bacterium *Wolbachia* (10,11). Oral administration of an iron chelator in the mosquito species *Anopheles albimanus* resulted in strain-specific differences in the severity of infections with the malaria parasite *Plasmodium berghei* there were fewer oocysts (parasites) on the midguts of one strain of *A. albimanus*, but the opposite result occurred in a different strain (12). Taken together, these studies indicate that the effect of dietary iron on infection outcome can depend on the specific host, the specific infectious microbe, and even the host strain.

The goal of this study was to determine whether dietary iron supplementation increases the severity of extracellular bacterial infections in the hemolymph of larval stage *Manduca sexta. We* predicted that iron would accumulate in the hemolymph of *M. sexta* larvae fed an iron supplemented diet. Our hypothesis was that the extra iron would promote proliferation of bacteria injected into the hemolymph, possibly leading to decreased host survival. Because this study is the first to focus on the effect of dietary iron on extracellular bacterial infections, we anticipated that the results would improve our understanding of the relationship between dietary iron and infections in insects. We also hoped the results would be useful for designing insect diets that reduce susceptibility to infection.

Our overall experimental approach was to feed *M. sexta* larvae an artificial diet with or without iron supplementation, inoculate larvae with bacteria, and measure the effects of iron supplementation on bacterial load and larval survival. For these studies, the control diet contained approximately 1.2 mM iron, and the iron-supplemented diet contained 11.2 mM iron. The amount of iron in the iron-supplement diet is similar to that used in the studies involving S. *poulsonii* and *Wolbachia* but less than the study involving *R. oryzae* (9–11). For comparison, the amount of iron in tobacco leaves, a natural diet of *M. sexta* larvae, has been observed in the range of 40-1000 mg/kg wet weight (assuming fresh tobacco leaves are 80-85% water), which corresponds to 0.7-17.9 mM iron (13,14). We typically culture *M. sexta* larvae on diet containing kanamycin and tetracycline to limit bacterial growth in the diet and, thus, decrease the variability of lab-induced immune responses (15); however, antibiotics can affect bacterial load and larval survival by inhibiting growth of inoculated bacteria and altering the microbiome of the midgut (16–18). For this reason, we used both antibiotics-free and antibiotics-added diet for this investigation.

We used two species of bacteria for this study. *Escherichia coli* was our choice of a nonpathogenic species because iron uptake mechanisms in *E. coli* are well-studied, mutant strains deficient in iron uptake are available, and it has been used it for previous studies of immunity in *M. sexta* (19–22). *Enterococcus faecalis* was our choice of an entomopathogen because multiple iron uptake mechanisms in *E. faecalis* have been identified, and it has been used previously for initiating infections in *M. sexta* larvae via inoculation (23,24).

The results of this study do not support the hypothesis that dietary iron supplementation increases the severity of extracellular bacterial infections in the hemolymph of larval stage *M. sexta*. Although iron supplementation increased the iron content of hemolymph, this increase did not result in an increase in bacterial load. Furthermore, the effect of iron supplementation on larval survival depended on the species of bacteria used and the presence or absence of antibiotics. Our results are consistent with previous studies that have shown a complex relationship between dietary iron and infection outcome.

## Materials and Methods

### Insect culture

Our *M. sexta* colony was maintained at 26 °C as described previously, including the use of a wheat germ-based larval diet containing two antibiotics (0.25 g/L kanamycin monosulfate and 0.25 g/L tetracycline hydrochloride) (15). Larvae used for experiments were fed either the standard wheat germ-based (control) diet with or without antibiotics, or iron-supplemented diet with or without antibiotics. We estimated that the control diet contained approximately 1.2 mM iron (mainly from two diet components: Wesson’s Salt Mixture and wheat germ). Iron-supplemented diet contained an additional 10 mM iron from adding 3.15 g ferric ammonium citrate (composed of 17.9% iron) per liter of diet.

The following procedure was used to rear larvae for experiments. Control diet was added to cups of eggs. Newly hatched larvae were placed on control or iron-supplemented diet, two per cup, for three days, and thereafter, larvae were reared individually as described previously (15). All comparisons between control and iron-treated larvae were made between larvae from the same batch of eggs. Regardless of diet type, a small percentage of larvae failed to thrive and were excluded from our analyses.

## Bacteria

### E. coli

Two strains of *E. coli* from the Keio Collection of single-gene knockouts were obtained from the Coli Genome Stock Center (20). Strain #7636 is the parent strain for the collection, and *we* refer to it as wild-type. Strain #11768 is identical to the parent strain except that it has a mutation in the entA gene (ΔentA734::kan); this mutation blocks the bacteria from synthesizing enterobactin, a siderophore needed for good growth in low iron environments (25). To prepare *E. coli* for inoculations, the following procedure was used. A fresh colony was added to 100 mL Luria-Bertani (LB) broth in a 500 mL flask, and the culture was grown at 37 °C with shaking for approximately 18 h. Bacteria were collected from 25 mL of culture by centrifugation at 2,612 x g for 15 minutes, washed by resuspending the bacteria in 0.85% NaCI, and centrifuged at 2,612 x g for 15 minutes. The bacterial pellet was resuspended in 0.85% NaCI, and the density was adjusted to an OD_600_ of 3, corresponding to approximately 1.2 × 10^6^ colony forming units (CFU) per μL. The expected CFU per μL was verified by plating dilutions on LB agar plates, growing colonies overnight at 37 °C, and counting the resulting colonies.

### E. faecalis

The *E. faecalis* strain designated OG1RF was obtained from the American Type Culture Collection. This strain was chosen because it is suitable for use in a biosafety level 1 lab and it has been used previously to infect *M. sexta* larvae (23). To prepare *E. faecalis* for inoculations, the following procedure was used. Unless otherwise noted, 100 mL Todd Hewitt (TH) broth in a 500 mL flask was inoculated with a fresh colony, and the culture was grown at 37 °C with shaking for approximately 26 h. Three experiments were performed with LB broth rather than TH, and these exceptions are noted in the relevant figure legend. Bacteria were collected by centrifugation at 2,612 x g for 15 minutes, washed by resuspending the bacteria in 0.85% NaCI, and centrifuged at 2,612 x g for 15 minutes. The bacterial pellet was resuspended in 0.85% NaCI, and the density was adjusted to an OD_600_ of 2, 0.2, or 0.02, corresponding to approximately 1.6 × 10^6^,1.6 × 10^5^, or 1.6 × 10^4^ colony forming units (CFU) per μL. The expected CFU per μL was verified by plating dilutions on TH agar plates, growing colonies overnight at 37 °C, and counting the resulting colonies.

Two experiments involved inoculation of larvae with dead *E. faecalis*. For the first experiment, the bacteria were grown and washed as usual, then resuspended in 0.37% formaldehyde in 0.85% NaCI, and placed on a rotator at room temperature for 3 days. The treated bacteria were washed three times and resuspended in 0.85% NaCI to an OD_600_ of 20, and then tested for viability. Because viability was not 0%, the formaldehyde-treated bacteria were heated to 70 °C for 1 h. To verify that the two-step killing process was effective, 50 μL bacteria were spread on an LB agar plate and incubated overnight at 37 °C. No colonies were observed. The killed bacteria were diluted 10 fold, corresponding to an OD_600_ of 2, before use. For the second experiment, bacteria were grown, washed, resuspended in 20 mL 0.37% formaldehyde in 0.85% NaCI, and placed on a rotator at room temperature for 7 days. The killed bacteria were washed three times with 10 mL 0.85% NaCI and resuspended in 4.5 mL 0.85% NaCI to correspond to an OD_600_ of 20. To verify that the bacteria were dead, 50 μL bacteria were spread on an LB agar plate and incubated overnight at 37 °C. No colonies were observed.

### Inoculations

Larvae fed antibiotics-free diet were inoculated when they were 11 days old, corresponding to one-day-old fifth instar larvae. Larvae fed antibiotics-containing diet developed more slowly, and so were inoculated at 12 days old, corresponding to a mix of one-day- and two-day-old fifth instar larvae. To anesthetize larvae, they were buried in ice for at least 8 minutes. Anesthetized larvae were injected with 50 μL bacteria with the use of a 1 mL syringe attached to a 30 gauge needle. Inoculated larvae were reared as usual.

### Measuring bacterial load

To estimate the number of live bacteria in hemolymph, we used a track dilution method that was based on previously published methods (26,27). Unless otherwise noted, this procedure was performed 24 h post-inoculation. First, larvae were chilled on ice for at least 5 minutes, and the area around the horn was cleaned with 70% ethanol. A cut was made at the base of the horn, and a few drops of hemolymph were collected in a sterile tube on ice. A 1:5 dilution was made by adding 20 μL hemolymph to 80 μL sterile 0.85% NaCI, and then up to five additional dilutions, each 10-fold, were made. The 1:5 dilutions of hemolymph contained antimicrobial activity that interfered with the assay; therefore, only the higher dilutions were used for data analysis. Diluted hemolymph was analyzed by placing 20 μL of each dilution at the top of square, gridded LB (for *E. coli)* or TH (for *E. faecalis)* agar plates, the plates were tilted until the tracks ran the length of the plate, and then the plates were incubated at 37 °C overnight. Colonies in tracks with suitable colony densities were counted, and data were converted to colony forming units (CPUs) per μL hemolymph.

### Monitoring larval survival

Feeding-stage larvae were observed each day, and any deaths were recorded. Once larvae stopped feeding and entered the wandering stage, they were placed in groups in boxes filled with vermiculite, and the number of larvae that reached the pupal stage was recorded. For experiments designed to generate survival curves, all larvae were monitored each day until they either died or pupated. Note that the larvae used for monitoring survival were not the same larvae as those used for measuring bacterial load because the wounding procedure used for collecting hemolymph can affect survival.

### Iron content assay

The iron content of larval hemolymph was measured by a FerroZine-based method that has been described previously (8) with several modifications. Larvae were fed control or iron-supplemented diet without antibiotics. Hemolymph was collected from four control and four iron-treated two-day-old fifth instar larvae by chilling them on ice, snipping off a proleg, and collecting hemolymph into chilled tubes. Hemolymph was stored frozen. After thawing, 300 μL hemolymph was mixed with 65 μL 12.1 N hydrochloric acid, and proteins were hydrolyzed at 95 °C for 20 minutes. Blanks were made by substituting hemolymph with deionized water. Samples were centrifuged for 16,000 x g for 5 min, and 200 μL supernatant was transferred to a new tube. To remove any remaining insoluble material, the samples were centrifuged for 16,000 x g for 2 min, and then 150 μL supernatant was mixed with 60 μL 75 mM ascorbic acid to reduce ferric ions to ferrous ions. After 10 min, the samples were mixed with 60 μL 10 mM FerroZine, and 120 μL 10.4 M ammonium acetate was added to increase the pH, allowing iron-FerroZine complex formation. To detect the iron-FerroZine complex, 300 μL of samples and blanks were added to wells of a 96 well plate, and absorbance was measured at 562 nm. The concentration of iron was calculated based on a molar extinction coefficient of the iron-FerroZine complex of 27,900 M ^−1^ cm^−1^ (28).

### Statistics

Statistical analyses were performed with GraphPad Prism version 5.04. An unpaired *t-* test with Welch’s correction was used to test for a statistically significant difference between iron concentrations in hemolymph. Contingency tables containing percent survival data were evaluated with a Fisher’s exact test. Differences in survival curves were evaluated with a log-rank test. Differences in the number of days to reach the wandering stage, which were based on non-continuous data, were tested with a Mann Whitney test. Data representing CFU per μL hemolymph were first tested for normality with a D’Agostino and Pearson omnibus normality test. For data that passed the normality test, differences in means were evaluated with an unpaired *t*-test, and means with standard deviations (SDs) are shown in graphs. For data that did not pass the normality test, differences in medians were evaluated with a Mann Whitney test, and medians with interquartile range (IQR) are shown in graphs. Samples sizes for all experiments are indicated in the figure legends. We did not perform statistical analyses of differences between experiments (for example, between experiments with and without antibiotics) because of uncontrolled variables.

## Results and Discussion

### Dietary iron supplementation increased the amount of iron in hemolymph

We predicted that dietary iron supplementation would increase the amount of iron in hemolymph. To test this hypothesis, we used a FerroZine-based assay to measure the iron content of control and iron-treated larvae. We found that hemolymph from larvae fed iron-supplemented diet had approximately 20 times more iron than hemolymph from control larvae (with a mean ± SD of 680 ± 91 μM for iron-treated larvae and 33 ± 13 μM for control larvae) (Fig 1). Assuming that at least some of the additional iron is accessible to bacteria, this finding supports our hypothesis that iron supplementation could increase the proliferation of bacteria in hemolymph, leading to an increase in infection severity.

**Fig 1.**
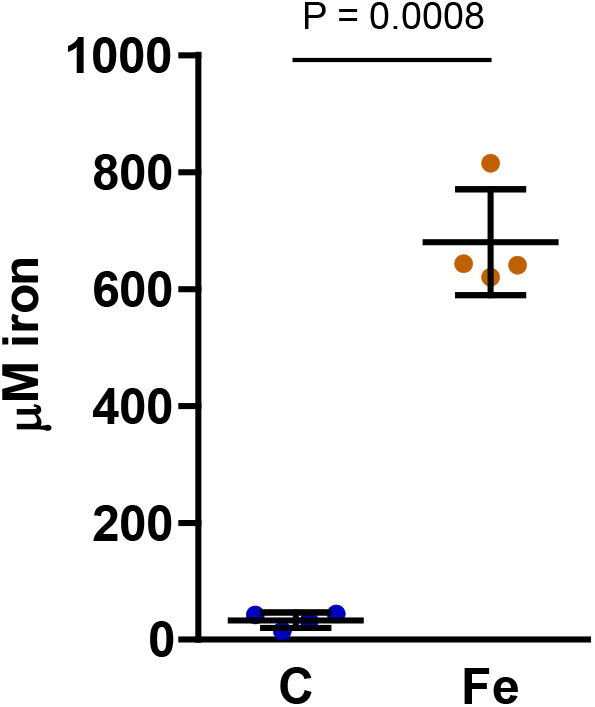
Iron supplementation increased iron content of hemolymph. Hemolymph was collected from fifth instar larvae that were reared on control (C) or iron-supplemented (Fe) diet without antibiotics, and a FerroZine-based assay was used to measure iron content. An unpaired *t*-test with Welch’s correction was used to test for statistical significance, and means ± SD are shown (n = 4).

### Dietary iron supplementation had little effect on naive larvae

To verify that the amount of iron in the iron-supplemented diet was not discernibly toxic to larvae, we fed larvae control or iron-supplemented diet, with or without antibiotics, and recorded growth rate, weight and survival. We found that in the absence of antibiotics, the iron-supplemented diet slowed the growth rate slightly, as demonstrated by the slightly longer time required for the iron-treated larvae to reach the wandering stage (Fig 2A); however, despite the slight developmental delay, the iron-treated larvae reached the same maximum weight as control larvae (Fig 2B). In the presence of antibiotics, neither growth rate nor maximum larval weight were affected by iron supplementation (Figs 2A and 2B). Regardless of the presence or absence of antibiotics, iron supplementation did not decrease larval survival (Fig 2C).

**Fig 2.**
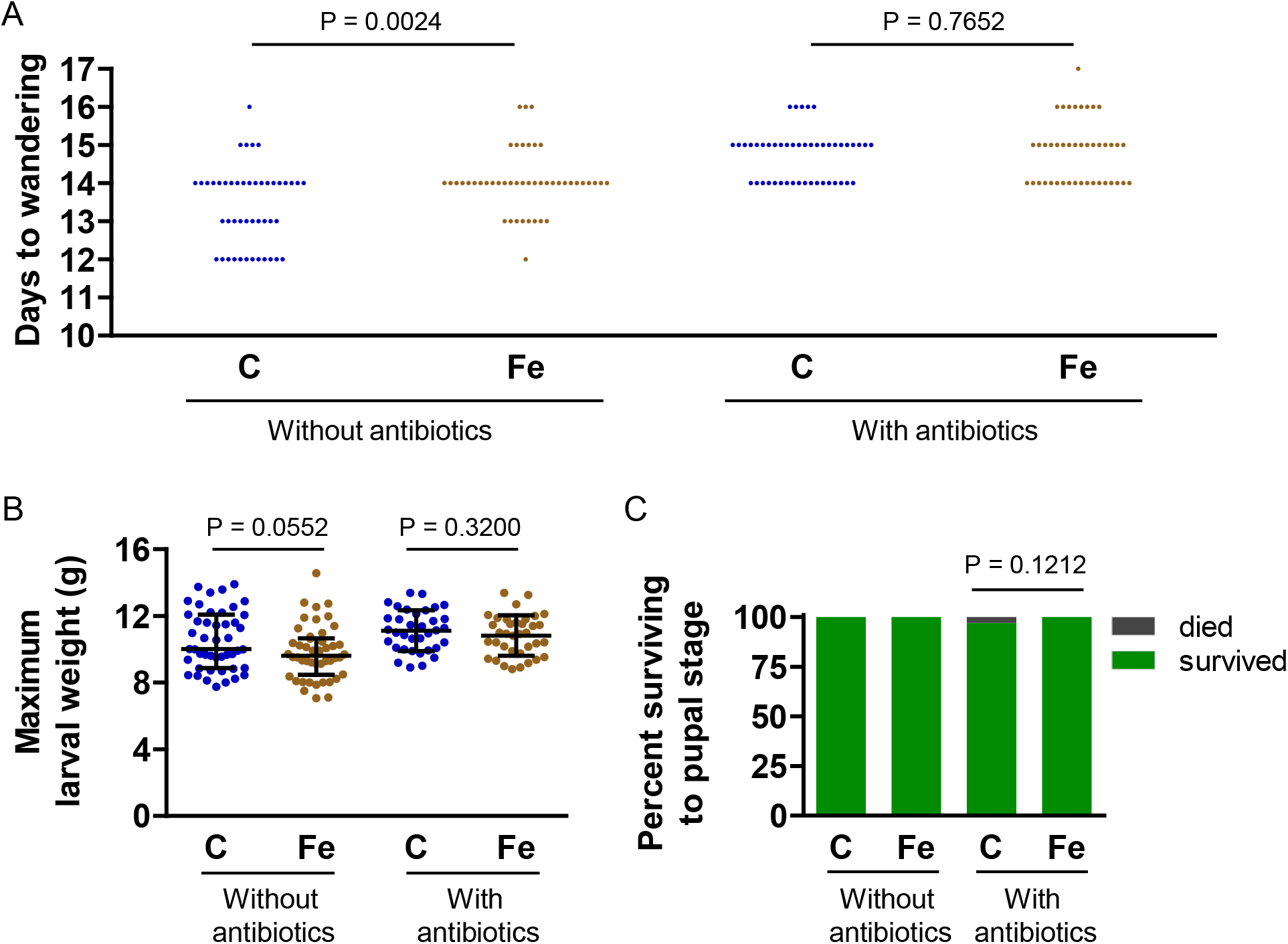
Iron supplementation had little effect on growth rate, weight and survival of naive larvae. Larvae were cultured on control (C) or iron-supplemented (Fe) diet, with or without antibiotics. A) Number of days from hatching to wandering stage. A Mann Whitney test was used to test for statistical significance (n = 43–47). B) Maximum larval weight. For the noantibiotics diet, a Mann Whitney test was used to test for statistical significance, and medians ± IQR are shown (n = 48). For the antibiotics-containing diet, an unpaired *t*-test was used to test for statistical significance, and means ± SD are shown (n = 36). C) Percentage of larvae surviving to the pupal stage. A Fisher’s exact test was used to test for statistical significance (without antibiotics, n = 48; with antibiotics, n = 36). For all experiments, the data presented were pooled from two or three biological replicates.

### Dietary iron supplementation did not affect the outcome of *E. coli* infections

To test the hypothesis that dietary iron supplementation increases the severity of *E. coli* infections, we inoculated larvae with *E. coli* at an OD_600_ of 3 and measured larval survival and number of bacteria in hemolymph. As expected, infection with nonpathogenic *E. coli* had no effect on larval survival (Fig 3A); however, contrary to our prediction, iron supplementation did not lead to an increase in the number of bacteria in hemolymph, regardless of whether the diet contained antibiotics (Fig 3B). At 24 hours post-inoculation, the number of bacteria in hemolymph was low, with a median CFU per μL ranging from 0 to 15. Assuming a larval hemolymph volume of roughly 1 mL, the number of live bacteria decreased from 6 × 10^7^ CFU per larva at the time of injection to a median of ≤ 1.5 × 10^4^ CFU per larva 24 hours later. At 48 hours post-inoculation, even fewer bacteria remained. These results suggest that most of the *E. coli* injected into larvae were killed by the larval immune system.

**Fig 3.**
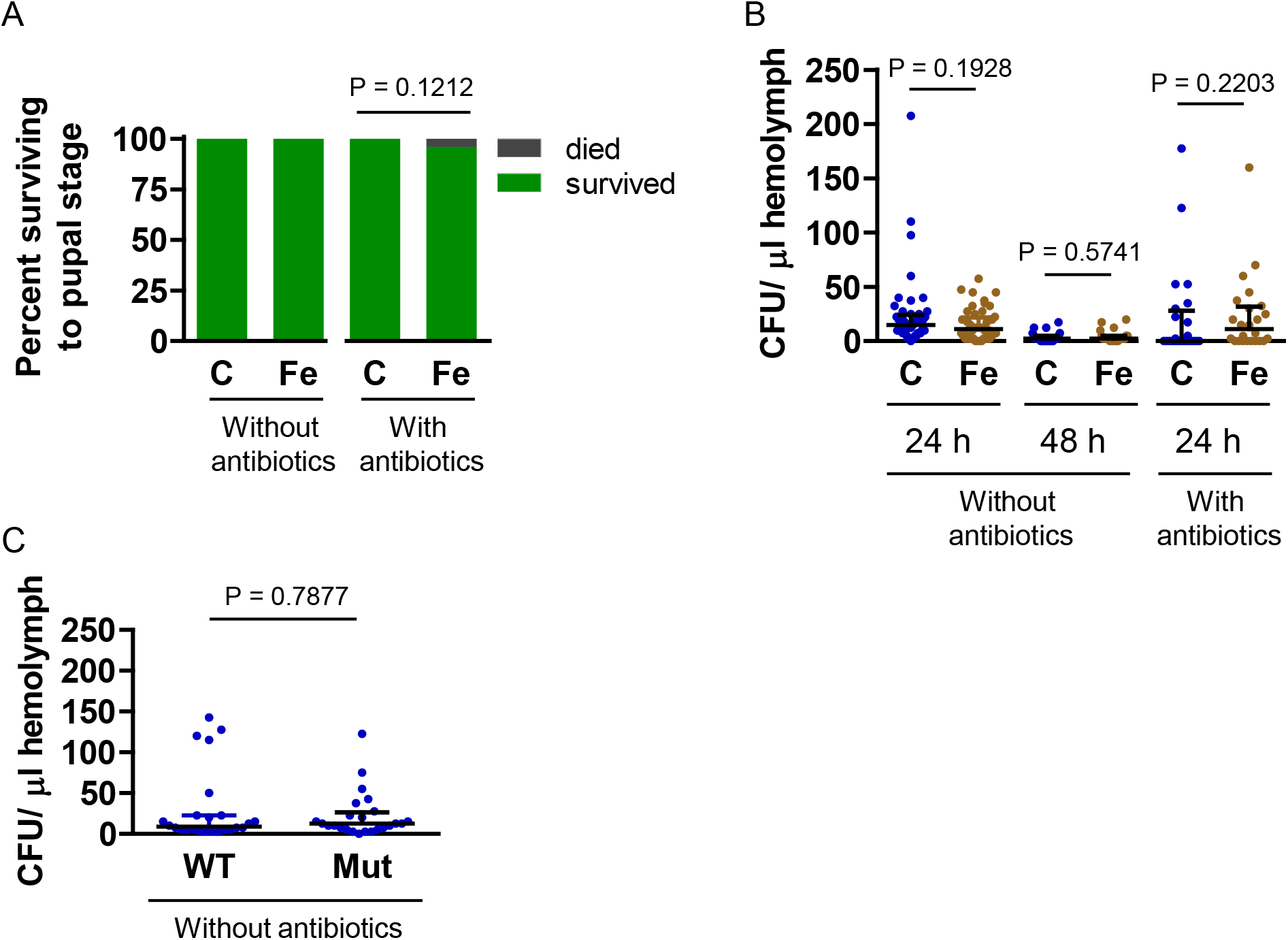
Iron supplementation had no effect on the severity of *E. coli* infections. Larvae cultured on control (C) or iron-supplemented (Fe) diet, with or without antibiotics, were inoculated with *E. coli* at an OD_600_ of 3, and infection severity was assessed. A) Percentage of inoculated larvae surviving to the pupal stage. A Fisher’s exact test was used to test for statistical significance (n = 24). B) The number of CFU per μL hemolymph. A Mann Whitney test was used to test for statistical significance; medians with upper IQR are shown (without antibiotics at 24 h, n = 48; the other two experiments, n = 24). The data for the first experiment in panel B are pooled from two biological replicates. C) Larvae fed the control diet without antibiotics were inoculated with either wild-type (WT) or a mutant (Mut) strain of *E. coli* that grows poorly in low iron conditions, and CFU per μL hemolymph 24 h after inoculation was determined. A Mann Whitney test was used to test for statistical significance; medians with upper IQR are shown (n = 24).

A possible explanation for why iron supplementation did not increase bacterial load is that *E. coli* in hemolymph may enter a quiescent state that does not require iron uptake. This could happen, for example, if larval hemolymph did not support the growth of *E. coli;* however, proliferation of *E. coli* in cell-free hemolymph in vitro suggests that hemolymph can support *E. coli* growth in vivo (29). A second possible explanation is that the amount of iron in control hemolymph may be sufficient for optimal bacterial growth. As an initial test of this hypothesis, we took advantage of a mutant strain of *E. coli* that does not synthesize the siderophore enterobactin and, thus, does not grow well in low iron environments (25). We inoculated control larvae with either wild-type or mutant *E. coli*, and then measured bacterial load at 24 hours post-inoculation. The number of mutant *E. coli* was the same as the number of wild-type *E. coli* (Fig 3C), suggesting that the amount of iron in hemolymph was not growth limiting.

### The effects of dietary iron supplementation on the outcome of *E. faecalis* infections was mixed

To test the hypothesis that iron supplementation would increase the severity of *E. faecalis* infections, we used four experimental conditions: antibiotics-free diet with *E*.*faecalis* at an OD_600_ of 0.02 or 0.2, and antibiotics-containing diet with *E*.*faecalis* at an OD_600_ of 0.2 or 2. Because *E*.*faecalis* is an entomopathogen (23,30), we anticipated some larval death for both diet treatment groups, but we predicted that the iron-supplemented diet would decrease survival by promoting bacterial proliferation.

As expected, when fed an antibiotics-free diet, control and iron-treated larvae were killed by *E. faecalis* infections. Whereas injection of *E. coli* at an OD_600_ of 3 was not lethal, injection of *E. faecalis* at an OD_600_ of 0.2 killed nearly all larvae within two days (Fig 4A). The number of *E. faecalis* increased from 8 × 10^6^ to 10^8^–10^9^ bacteria per larva in the 24 hours after inoculation, indicating that larval hemolymph can support the growth of *E. faecalis* (Fig 4B). However, contrary to our hypothesis, we observed no significant difference in survival or number of bacteria between control and iron-treated larvae (Figs 4A and 4B).

**Fig 4.**
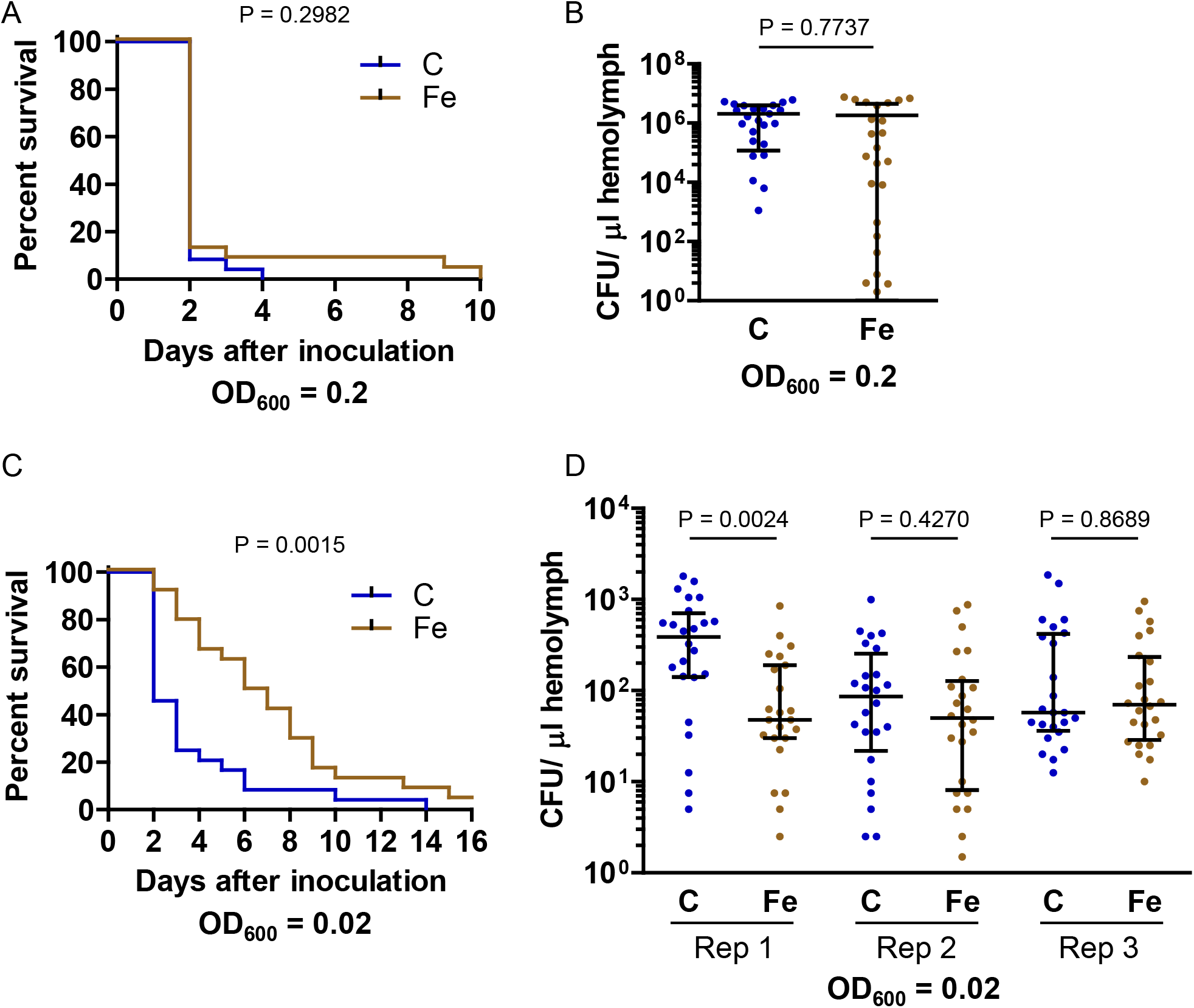
Without antibiotics, iron supplementation prolonged survival of larvae infected with *E. faecalis* without decreasing bacterial load. Larvae cultured on control (C) or iron-supplemented (Fe) diet without antibiotics were inoculated with *E. faecalis*. A) Larvae were inoculated with *E. faecalis* at an OD_600_ of 0.2, and mortality was monitored. The resulting survival curves were compared by performing a log-rank test (n = 24). B) Larvae were inoculated with *E. faecalis* at an OD_600_ of 0.2, and CFU per μL hemolymph was measured after 24 h. An unpaired *t*-test was used to test for statistical significance, and means ± SD are shown (n = 24). C) Larvae were inoculated with *E. faecalis* at an OD_600_ of 0.02, and mortality was monitored. The resulting survival curves were compared by performing a log-rank test (n = 24). D) Larvae were inoculated with *E. faecalis* at an OD_600_ of 0.02, and CFU per μL hemolymph was measured after 24 h. The results of three biological replicates are presented. A Mann Whitney test was used to test for statistical significance; medians ± IQR are shown (n = 24).

By injecting *E. faecalis* at an OD_600_ of 0.02, we were able to avoid the rapid, severe infections produced by injecting *E. faecalis* at an OD_600_ of 0.2. Under these conditions, we saw a difference in survival, but, surprisingly, instead of decreasing larval survival, iron supplementation extended survival time (Fig 4C and S1 Fig). These data suggest that dietary iron supplementation may temporarily protect larvae from infection with *E. faecalis*. To determine whether the longer survival of iron-treated larvae corresponded to a reduced bacterial load, we measured the number of bacteria in hemolymph 24 hours after inoculation. For one biological replicate, iron-treated larvae had fewer bacteria, but for the other two replicates there was no significant difference between diet treatments (Fig 4D). These results suggest that the cause of the extended survival of iron-treated larvae is unrelated to bacterial load. The finding that iron supplementation may temporarily protect larvae from *E. faecalis*-induced death was unexpected. Given that dietary iron supplementation can affect the gut microbiome of humans (31,32), a potential explanation is that iron supplementation may affect the gut microbiome in a way that influences infection tolerance (17).

Because our standard larval diet includes kanamycin and tetracycline, we wanted to know how iron supplementation in the presence of these antibiotics affects infection severity. We found that the addition of antibiotics considerably altered the effects of iron supplementation. As described above, when injected with *E*.*faecalis* at an OD_600_ of 0.2, most larvae fed antibiotics-free diet developed a high bacterial load and died (Fig 4A); in contrast, larvae fed the antibiotics-containing diet survived the infection and had a very low bacterial load, with no significant difference between control and iron-treated larvae (Figs 5A and 5B). These results suggest that antibiotics can be transported from the diet to the larval hemolymph where they can kill *E. faecalis* (a phenomenon we did not observe with *E. coli)*.

**Fig 5.**
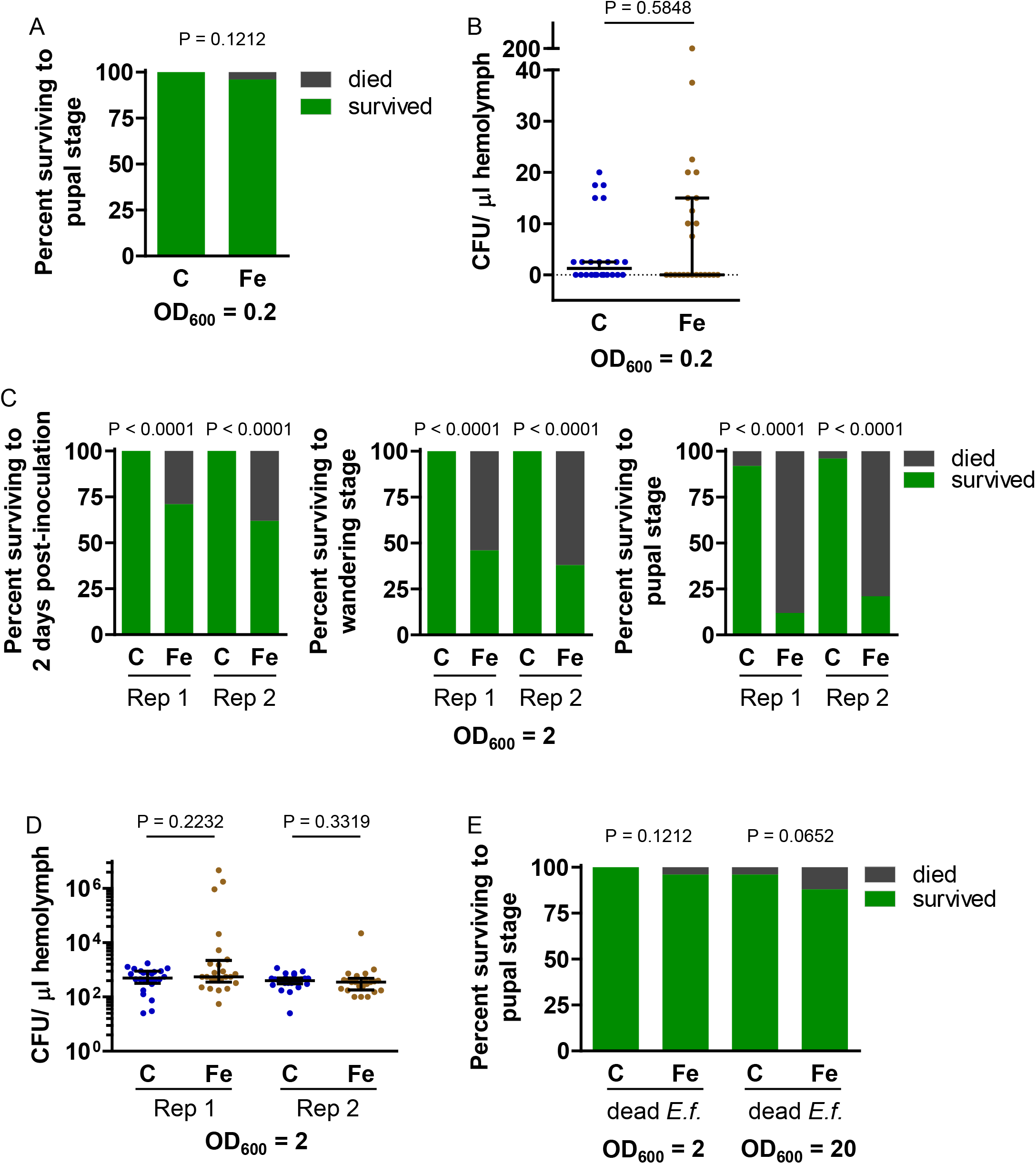
With antibiotics, iron supplementation decreased survival of larvae infected with *E. faecalis* without increasing bacterial load. Larvae cultured on control (C) or iron-supplemented (Fe) antibiotics-containing diet were inoculated with *E*.*faecalis*. A) Larvae were inoculated with *E*.*faecalis* at an OD_600_ of 0.2, and survival to the pupal stage was monitored. A Fisher’s exact test was used to test for statistical significance (n = 24). B) Larvae were inoculated with *E. faecalis* at an OD_600_ of 0.2, and CFU per μL hemolymph was measured after 24 h. A Mann Whitney test was used to test for statistical significance; medians with upper IQR are shown (n = 24). C) Larvae were inoculated with *E. faecalis* at an OD_600_ of 2, and mortality was monitored. The percentage surviving at two days post-inoculation, the wandering stage, and the pupal stage are shown. The results of two biological replicates are presented. A Fisher’s exact test was used to test for statistical significance (n = 24). D) Larvae were inoculated with *E. faecalis* at an OD_600_ of *2*, and CFU per μL hemolymph was measured after 24 h. The results from two biological replicates are shown. A Mann Whitney test was used to test for statistical significance; medians ± IQR are shown (n = 24). E) Larvae were inoculated with dead *E. faecalis* at an OD_600_ of 2 or 20, and survival to the pupal stage was monitored. A Fisher’s exact test was used to test for statistical significance (n = 24). *E. faecalis* was cultured in LB broth for experiments presented in A, B, and C replicate 1, whereas *E. faecalis* was cultured in TH broth for experiments presented in C replicate 2 and D.

For the next set of experiments, we fed larvae antibiotics-containing diet and injected *E. faecalis* at an OD_600_ of 2. Under these experimental conditions, the iron-treated larvae were significantly more likely than control larvae to die; this effect was significant at 2 days postinoculation and was even more dramatic by the end of the larval period (Fig 5C). This result supported our hypothesis that iron supplementation may increase infection severity; however, surprisingly, decreased survival did not correspond to an increased bacterial load (Fig 5D). Excess iron, immune reactions, and antibiotics (including tetracyclines) can all cause physiological stress (33–35), and it appears that the inoculated larvae that were exposed to antibiotics and extra iron may have died from a lethal combination of these stresses. We wondered whether live bacteria were necessary for the high mortality or if an infection-free immune response (36,37) would be sufficient. To answer this question, we inoculated larvae with killed *E. faecalis* at an OD_600_ of 2 and then monitored larval survival. We found that killed bacteria did not significantly decrease the survival of iron-treated larvae (Fig 5E). Even a ten fold increase in dead bacteria did not result in a significant decrease in survival (Fig 5E). These results suggest that live *E. faecalis* were needed for high larval mortality. Our experiments involving the injection of a large number of *E. faecalis* into larvae fed antibiotics-containing diet demonstrate that iron supplementation can increase larval mortality, but this outcome is independent of bacterial load, and the mechanism underlying the phenomenon is unknown.

## Conclusions

The central hypothesis of this study was that dietary iron supplementation would increase the amount of iron in larval hemolymph, promoting the growth of infectious microbes, possibly resulting in larval death. Although we verified that iron supplementation increased the iron content of hemolymph, we saw no evidence that additional iron increased proliferation of either *E. coli* or *E*.*faecalis* under any of our experimental conditions. Different bacterial species or different experimental conditions (e.g., more or less iron in the diet) could produce different outcomes; however, the results of this study support the conclusion that dietary iron supplementation does not promote bacterial proliferation in *M. sexta* hemolymph.

Contrary to our model, there were differences in survival between control and iron-treated larvae that were independent of the number of bacteria in the hemolymph. Surprisingly, when larvae were injected with *E. faecalis*, the effect of iron supplementation on survival depended on whether the diets contained antibiotics. In the absence of antibiotics, iron-treated larvae survived longer than control larvae, whereas, in the presence of antibiotics, iron-treated larvae were less likely to survive than control larvae. Given that antibiotics in the diet probably alter the gut microbiome, changes in the microbiome may underlie the difference in outcomes. But less obvious explanations are also possible. For example, the effect of antibiotics on the survival of iron-treated larvae may be mediated by chemical interactions between iron and the tetracycline in the diet (38,39). More studies are needed to explore these possibilities.

The previously published research on the effects of dietary iron on immunity suggest that the relationship between dietary iron and infection outcome is complex (5,9–12). The results of our study support this viewpoint.

## Acknowledgments

We are grateful to Kristin Michel, Dewey Leierer, and Shirley Luckhart for helpful comments regarding this work. We thank Bianca Morejon for advice about culturing *E. faecalis* and counting bacteria, and Lisa Brummett for information about working with different strains of *E. coli*.

## Funding

The research reported in this publication was supported by the National Institute of General Medical Sciences of the National Institutes of Health (NIH) under Award Number R35GM141859 (MK), and by the intramural research program of the U.S. Department of Agriculture, National Institute of Food and Agriculture, Hatch project 7003374 (MK). The content is solely the responsibility of the authors and does not necessarily represent the official views of the NIH. The findings and conclusions in this publication have not been formally disseminated by the U. S. Department of Agriculture and should not be construed to represent any agency determination or policy. This is contribution 25-236-J from the Kansas Agricultural Experiment Station. The funders had no role in study design, data collection and analysis, decision to publish, or preparation of the manuscript.

## Supporting information

**S1 Fig.In the absence of antibiotics, iron supplementation decreased *E. faecalis-induced* mortality of feeding stage larvae.**Larvae cultured on control (C) or iron-supplemented (Fe) diet without antibiotics were inoculated with *E*.*faecalis* at an OD_600_ of 0.02, and mortality was monitored. The percentage of larvae surviving at three days post-inoculation, the wandering stage, and the pupal stage are shown. A Fisher’s exact test was used to test for statistical significance (n = 24).

## Notes

### Competing Interest Statement

The authors have declared no competing interest.

### Summary of Updates

Added funding information, including required statements from funding agencies, to the text.

## References

1. Gorman MJ. Iron Homeostasis in Insects. Annu Rev Entomol. 2023 Jan 23;68:51–67.

2. Andrews SC, Robinson AK, Rodríguez-Quiñones F. Bacterial iron homeostasis. FEMS Microbiol Rev. 2003 Jun 1;27(2–3):215–37.

3. Weinberg ED. Iron availability and infection. Biochim Biophys Acta BBA - Gen Subj. 2009 Jul;1790(7):600–5.

4. Ong ST, Ho JZS, Ho B, Ding JL. Iron-withholding strategy in innate immunity. Immunobiology. 2006;211(4):295–314.

5. Hrdina A, Iatsenko I. The roles of metals in insect-microbe interactions and immunity. Curr Opin Insect Sci. 2021 Dec 21;49:71–7.

6. Brownlie JC, Cass BN, Riegler M, Witsenburg JJ, Iturbe-Ormaetxe I, McGraw EA, et al. Evidence for Metabolic Provisioning by a Common Invertebrate Endosymbiont, Wolbachia pipientis, during Periods of Nutritional Stress. PLOS Pathog. 2009 Apr 3;5(4):e1000368.

7. Peng Z, Dittmer NT, Lang M, Brummett LM, Braun CL, Davis LC, et al. Multicopper oxidase-1 orthologs from diverse insect species have ascorbate oxidase activity. Insect Biochem Mol Biol. 2015 Apr;59:58–71.

8. Missirlis F, Holmberg S, Georgieva T, Dunkov BC, Rouault TA, Law JH. Characterization of mitochondrial ferritin in Drosophila. Proc Natl Acad Sci U S A. 2006 Apr 11;103(15):5893–8.

9. Chamilos G, Lewis RE, Hu J, Xiao L, Zal T, Gilliet M, et al. Drosophila melanogaster as a model host to dissect the immunopathogenesis of zygomycosis. Proc Natl Acad Sci U S A. 2008 Jul 8;105(27):9367–72.

10. Marra A, Masson F, Lemaitre B. The iron transporter Transferrin 1 mediates homeostasis of the endosymbiotic relationship between Drosophila melanogaster and Spiroplasma poulsonii. microLife. 2021 Jan 1;2:uqab008.

11. Currin-Ross D, Husdell L, Pierens GK, Mok NE, O’Neill SL, Schirra HJ, et al. The Metabolic Response to Infection With Wolbachia Implicates the Insulin/Insulin-Like-Growth Factor and Hypoxia Signaling Pathways in Drosophila melanogaster. Front Ecol Evol [Internet]. 2021 Mar 22 [cited 2026 Jan 19];9. Available from: https://www.frontiersin.org/journals/ecology-and-evolution/articles/10.3389/fevo.2021.623561/full

12. Maya-Maldonado K, Cardoso-Jaime V, González-Olvera G, Osorio B, Recio-Tótoro B, Manrique-Saide P, et al. Mosquito metallomics reveal copper and iron as critical factors for Plasmodium infection. PLoS Negl Trop Dis. 2021 Jun 23;15(6):e0009509.

13. Li Y, Liu F, Sun S, Xiang Y, Jiang X, He J. Metabolome of flue-cured tobacco is significantly affected by the presence of leaf stem. BMC Plant Biol. 2023 Feb 13;23(1):89.

14. Dospatliev L, Zaprjanova P, Ivanov K, Angelova V. Correlation between soil characteristics and iron content in aboveground biomass of Virginia tobacco. Bulg J Agric Sci. 2014;20(6):1380–5.

15. Kanost M, Gorman M, Rowe J, Brummett L. Protocols for maintaining a colony of Manduca sexta (tobacco hornworm) suitable for studies of innate immunity and other investigations of a model insect species [Internet]. OSF; 2025 [cited 2025 May 12]. Available from: https://osf.io/zduy8_v1

16. Hammer TJ, Janzen DH, Hallwachs W, Jaffe SP, Fierer N. Caterpillars lack a resident gut microbiome. Proc Natl Acad Sci USA. 2017 Sep 5;114(36):9641–6.

17. Lesperance DN, Broderick NA. Microbiomes as modulators of Drosophila melanogaster homeostasis and disease. Curr Opin Insect Sci. 2020 Jun 1;39:84–90.

18. Grenier T, Leulier F. How commensal microbes shape the physiology of Drosophila melanogaster. Curr Opin Insect Sci. 2020 Oct;41:92–9.

19. Braun V, Braun M. Iron transport and signaling in Escherichia coli. FEBS Lett. 2002 Oct 2;529(1):78–85.

20. Baba T, Ara T, Hasegawa M, Takai Y, Okumura Y, Baba M, et al. Construction of Escherichia coli K-12 in-frame, single-gene knockout mutants: the Keio collection. Mol Syst Biol. 2006 Jan;2(1):2006.0008.

21. Dunn PE, Drake DR. Fate of bacteria injected into naive and immunized larvae of the tobacco hornworm Manduca sexta. J Invertebr Pathol. 1983 Jan 1;41(1):77–85.

22. Booth K, Cambron L, Fisher N, Greenlee KJ. Immune Defense Varies within an Instar in the Tobacco Hornworm, Manduca sexta. Physiol Biochem Zool PBZ. 2015;88(2):226–36.

23. Mason KL, Stepien TA, Blum JE, Holt JF, Labbe NH, Rush JS, et al. From Commensal to Pathogen: Translocation of Enterococcus faecalis from the Midgut to the Hemocoel of Manduca sexta. mBio. 2011 May 17;2(3):10.1128/mbio.00065-11.

24. Brunson DN, Colomer-Winter C, Lam LN, Lemos JA. Identification of Multiple Iron Uptake Mechanisms in Enterococcus faecalis and Their Relationship to Virulence. Infect Immun. 2023 Apr 18;91(4):e0049622.

25. Young IG, Langman L, Luke RKJ, Gibson F. Biosynthesis of the Iron-Transport Compound Enterochelin: Mutants of Escherichia coli Unable to Synthesize 2,3-Dihydroxybenzoate. J Bacteriol. 1971 Apr;106(1):51–7.

26. Jett BD, Hatter, Kenneth L., Huycke, Mark M., and Gilmore MS. Simplified Agar Plate Method for Quantifying Viable Bacteria. BioTechniques. 1997 Oct 1;23(4):648–50.

27. Morejon B, Michel K. The expanded immunoregulatory protease network in mosquitoes is governed by gene coexpression. Proc Natl Acad Sci U S A. 2025 May 6;122(18):e2425863122.

28. Stookey LL. Ferrozine—a new spectrophotometric reagent for iron. Anal Chem. 1970 Jun 1;42(7):779–81.

29. Horohov DW, Dunn PE. Changes in the circulating hemocyte population of Manduca sexta larvae following injection of bacteria. J Invertebr Pathol. 1982 Nov 1;40(3):327–39.

30. Garsin DA, Frank KL, Silanpää J, Ausubel FM, Hartke A, Shankar N, et al. Pathogenesis and Models of Enterococcal Infection. In: Gilmore MS, Clewell DB, Ike Y, Shankar N, editors. Enterococci: From Commensals to Leading Causes of Drug Resistant Infection [Internet]. Boston: Massachusetts Eye and Ear Infirmary; 2014 [cited 2025 May 29]. Available from: http://www.ncbi.nlm.nih.gov/books/NBK190426/

31. Iddrisu I, Monteagudo-Mera A, Pověda C, Shahzad M, Walton GE, Andrews SC. A review of the effect of iron supplementation on the gut microbiota of children in developing countries and the impact of prebiotics. Nutr Res Rev. 2025 Jun;38(1):229–37.

32. Finlayson-Trick E, Nearing J, Fischer JAJ, Ma Y, Wang S, Krouen H, et al. The Effect of Oral Iron Supplementation on Gut Microbial Composition: a Secondary Analysis of a Double-Blind, Randomized Controlled Trial among Cambodian Women of Reproductive Age. Microbiol Spectr. 2023 May 18;11(3):e05273–22.

33. Korsloot A, van Gestel CAM, van Straalen NM. Environmental Stress and Cellular Response in Arthropods [Internet]. Boca Raton, Florida: CRC Press; 2004 [cited 2022 May 6]. Available from: https://www.journals.uchicago.edu/doi/full/10.1086/513335

34. Wang X, Ryu D, Houtkooper RH, Auwerx J. Antibiotic use and abuse: A threat to mitochondria and chloroplasts with impact on research, health, and environment. BioEssays. 2015;37(10):1045–53.

35. Adamo SA. The stress response and immune system share, borrow, and reconfigure their physiological network elements: Evidence from the insects. Horm Behav. 2017 Feb 1;88:25–30.

36. Hixson B, Huot L, Morejon B, Yang X, Nagy P, Michel K, et al. The transcriptional response in mosquitoes distinguishes between fungi and bacteria but not Gram types. BMC Genomics. 2024 Apr 9;25(1):353.

37. Kanost MR, Dai W, Dunn PE. Peptidoglycan fragments elicit antibacterial protein synthesis in larvae of Manduca sexta. Arch Insect Biochem Physiol. 1988;8(3):147–64.

38. Neuvonen PJ. Interactions with the absorption of tetracyclines. Drugs. 1976;11(1):45–54.

39. Wang H, Yao H, Sun P, Li D, Huang CH. Transformation of Tetracycline Antibiotics and Fe(II) and Fe(III) Species Induced by Their Complexation. Environ Sci Technol. 2016 Jan 5;50(1):145–53.

