## Supplementary figures and images for "The effects of dietary iron supplementation on bacterial infections in *Manduca sexta* larval hemolymph"

### Supplemental Fig 1

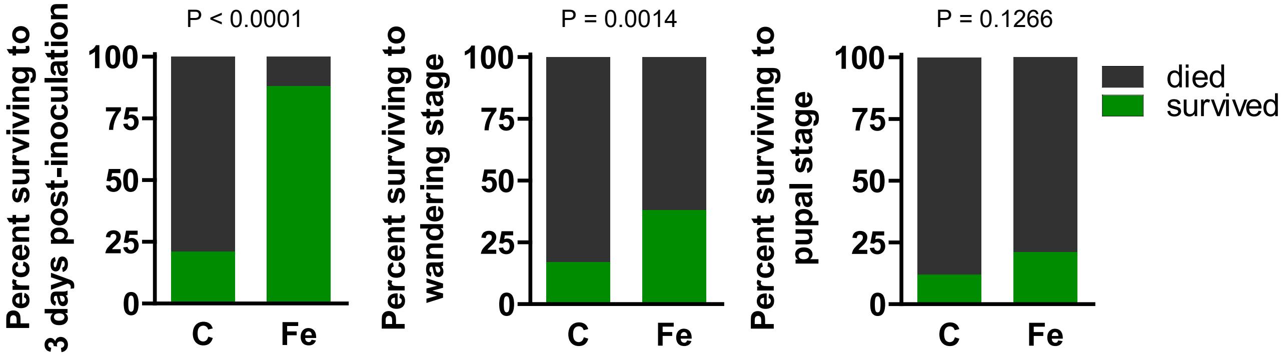
